# The Herpes Simplex Virus Tegument Protein pUL21 is Required for Viral Genome Retention Within Capsids

**DOI:** 10.1101/2022.07.14.500155

**Authors:** Ethan C.M. Thomas, Maike Bossert, Bruce W. Banfield

**Affiliations:** Department of Biomedical and Molecular Sciences, Queen’s University, Kingston, ON, Canada, K7L 3N6

**Author notes:** Corresponding Author: Bruce W. Banfield, Ph.D. Room 741 Botterell Hall, 18 Stuart Street Kingston, ON, Canada, K7L 3N6.

## Abstract

During virion morphogenesis herpes simplex virus nucleocapsids transit from the nucleoplasm to the cytoplasm, through a process called nuclear egress, where the final stages of virion assembly occur. Coupled to nuclear egress is a poorly understood quality-control mechanism that preferentially selects genome-containing C-capsids, rather than A- and B-capsids that lack genomes, for transit to the cytoplasm. We and others have reported that cells infected with HSV strains deleted for the tegument protein pUL21 accumulate both empty A-capsids and C-capsids in the cytoplasm of infected cells. Quantitative microscopy experiments indicated that C-capsids were preferentially selected for envelopment at the inner nuclear membrane and that nuclear integrity remained intact in cells infected with pUL21 mutants, prompting alternative explanations for the accumulation of A-capsids in the cytoplasm. More A-capsids were also found in the nuclei of cells infected with pUL21 mutants compared to their wild type (WT) counterparts, suggesting pUL21 might be required for optimal genome packaging or genome retention within capsids. In support of this, more viral genomes were prematurely released into the cytoplasm during pUL21 mutant infection compared to WT infection and led to enhanced activation of cellular cytoplasmic DNA sensors. Mass spectrometry and western blot analysis of WT and pUL21 mutant capsids revealed an increased association of the known pUL21 binding protein, pUL16, with pUL21 mutant capsids, suggesting that premature and/or enhanced association of pUL16 with capsids might result in capsid destabilization. Further supporting this idea, deletion of pUL16 from a pUL21 mutant strain rescued genome retention within capsids. Taken together, these findings suggest that pUL21 regulates pUL16 addition to nuclear capsids and that premature, and/or, over-addition of pUL16 impairs HSV genome retention within capsids.

**Importance:** pUL21 is a conserved, multifunctional alphaherpesvirus protein involved in cell-to-cell spread of infection, transport of capsids along microtubules, and regulation of the phosphorylation status of both viral and host proteins. pUL21 thereby controls diverse processes such as nuclear egress of capsids and cellular lipid trafficking. This study provides additional insight into HSV-1 and HSV-2 pUL21 activities and suggests that, by binding pUL16, pUL21 prevents pUL16 from interacting prematurely with nuclear capsids that would otherwise lead to capsid destabilization and premature ejection of viral genomes. Thus, prevention of pUL21 interaction with pUL16 may prove to be a useful strategy for interfering with virion assembly.

## Introduction

The alphaherpesviruses herpes simplex virus (HSV) 1 and 2 are highly prevalent human pathogens (1, 2). Viral genome synthesis, capsid assembly, and genome packaging into capsids occur in the nuclei of infected cells. Capsid assembly begins with the formation of a spherical two-shelled structure called a procapsid (3). The outer shell is comprised of 150 pentons and 11 hexons of the major capsid protein VP5, triplexes composed of proteins VP23 and VP19c that link the hexons and pentons, as well as one capsid portal composed of a dodecamer of pUL6, through which nascent genomes are packaged (3). The inner shell is the capsid scaffold and is constructed from the large and small scaffolding proteins pUL26 and pUL26.5. Genome packaging is initiated by the association of the terminase complex with the portal triggering a signal transduction event that leads to the activation of pUL26 protease activity and subsequent release of the scaffold. This results in angularization of the capsid shell as the genome is packaged into the capsid. Successful genome packaging results in a genome-containing capsid referred to as a C-capsid. In addition to C-capsids, two by-products of capsid maturation, A- and B-capsids, are also produced in infected nuclei. B-capsids are thought to have initiated proteolytic cleavage of the scaffolding proteins and become angularized without engaging the terminase complex (4, 5). As such, B-capsids are terminal products that lack genomic DNA and contain a cleaved form of the small scaffolding protein, VP22a, trapped inside the capsid. A-capsids arise through unsuccessful genome packaging events or through the inability of the capsid to retain the viral genome (4-6).Regardless of their origin, A-capsids are angularized and empty with no genomic DNA or scaffolding proteins present within their interior. In general, A-capsids are less abundant in infected cells compared to C-capsids as several mechanisms are utilized to retain the viral genome within capsids after packaging (3).

One of the most studied contributors to the retention of viral genomes within capsids are the capsid vertex specific component (CVSC) proteins pUL17, pUL25, and pUL36 (7-12). These proteins are bound to triplexes adjacent to capsid vertices and are required for stabilizing the capsid structure against the high internal pressure exerted by the packaged genome (7, 12-15). While CVSC proteins are found on A-, B-, and C-capsids, they are more abundant on C-capsids (14, 16). pUL17 is important for genome packaging and the efficient recruitment of pUL25 and pUL36 to capsids (12, 14). pUL25 on capsid vertices prevents the capsid structure from rupturing after packaging (17, 18). Further, pUL25 adjacent to the capsid portal is thought to extend over the portal, thereby plugging it after genome packaging (14). pUL36 is important for the recruitment of the tegument to capsids during maturation and for proper portal plug conformation (9, 14, 19). In addition to the CVSC proteins, capsids possess disulfide bonds that covalently link capsid proteins (20-22). Impairing these complexes results in the dissociation of pentons, triplexes, and CVSC proteins from the capsid and the loss of genomes from the capsid. Ultimately, the successful packaging and retention of the genome into a newly assembled capsid is only one hurdle HSV capsids must face in the virion maturation process.

After genome packaging, C-capsids must transit from the nucleus to the cytoplasm for the final stages of virion assembly (23). However, capsids are too large to pass through nuclear pore complexes. Thus, capsids undergo a specialized process conserved throughout the *Herpesviridae* family, called nuclear egress (23, 24). Nuclear egress is a complex and highly regulated process whereby C-capsids bud into the perinuclear space, acquiring a primary envelope derived from the inner nuclear membrane, followed by fusion of this envelope with the outer nuclear membrane, thereby releasing the capsid into the cytoplasm. Nuclear egress is also coupled to a poorly understood quality-control mechanism that preferentially selects C-capsids, rather than A- and B-capsids, to undergo primary envelopment at the inner nuclear membrane (25).

The capsid-associated tegument protein pUL21 is one of several viral proteins that function in nuclear egress and is conserved amongst the alphaherpesvirus subfamily (26-29). pUL21 forms a tripartite complex in the mature HSV virion, interacting with tegument protein pUL16, which in turn interacts with pUL11 (30-34). During the early stages of infection, pUL21 has been implicated in the trafficking of capsids to nuclei in HSV-1, HSV-2, and pseudorabies virus (PRV) infected cells as well as being required for optimal viral gene expression (28, 35-37). Late in infection, pUL21 regulates nuclear egress, tegument formation, cell-to-cell spread of infection and actives the ceramide transport protein, CERT (26, 27, 29, 38-40). Despite significant research into several pUL21 activities, the function pUL21 performs on capsids has yet to be thoroughly investigated. Previous studies from this laboratory and others have demonstrated that HSV-1 and HSV-2 pUL21 mutant strains accumulate both A- and C-capsids in the cytoplasm of infected cells (40, 41). In wild type (WT) virus infected cells, the majority of capsids found in the cytoplasm are C-capsids while A-capsids are almost exclusively found in the nucleus (3, 16). Currently, there is no explanation for the accumulation of A-capsids in the cytoplasm of cells infected with pUL21 deficient strains.

At least three hypotheses might explain the cytoplasmic accumulation of A-capsids in cells infected with pUL21 mutants. First, nuclear integrity may be disrupted in cells infected with pUL21 mutant strains leading to indiscriminate leakage of capsids from the nucleoplasm into the cytoplasm. Second, pUL21 may be required for the preferential selection of C-capsids for nuclear egress and its deletion may result in the selection of A-capsids for nuclear egress. Finally, it may be that viral genomes are not stably packaged into pUL21 mutant capsids resulting in premature expulsion of the genome from capsids once they reach the cytoplasm. The goal of this study was to evaluate these three hypotheses.

## Results

### Deletion of UL21 results in an increased abundance of nuclear and cytoplasmic A-capsids in infected cells

HSV-1 and HSV-2 pUL21 mutants (Δ21) accumulate both A- and C-capsids in the cytoplasm of infected cells (40, 41). Although this phenotype has been documented, quantification of A-capsids in HSV-1 and HSV-2 Δ21 infected cells has not been conducted.

To quantify the numbers of nuclear A-capsids in Δ21 infected cells, multiple WT and Δ21 mutant strains derived from HSV-1 and HSV-2 were used to infect Vero cells for transmission electron microscopy (TEM) analysis (Fig 1A-D). Cells infected with HSV-1 and HSV-2 Δ21 mutants had more nuclear A-capsids compared to WT infected cells (Fig 1E). Unpaired t-tests were conducted for HSV-2 (186, SD90e, HG52) and HSV-1 (F, KOS) strains to compare the percentage of nuclear A-capsids in Δ21 infections to WT infections. The Δ21 mutants of HSV-2 186 and SD90e, as well as HSV-1 F and KOS, had significantly more nuclear A-capsids present in infected cells compared to corresponding WT infections. There was no significant difference (p = 0.0846) between the HSV-2 HG52 Δ21 mutant and WT virus, despite the presence of more nuclear A-capsids in Δ21 infected cells.

**Fig 1.**
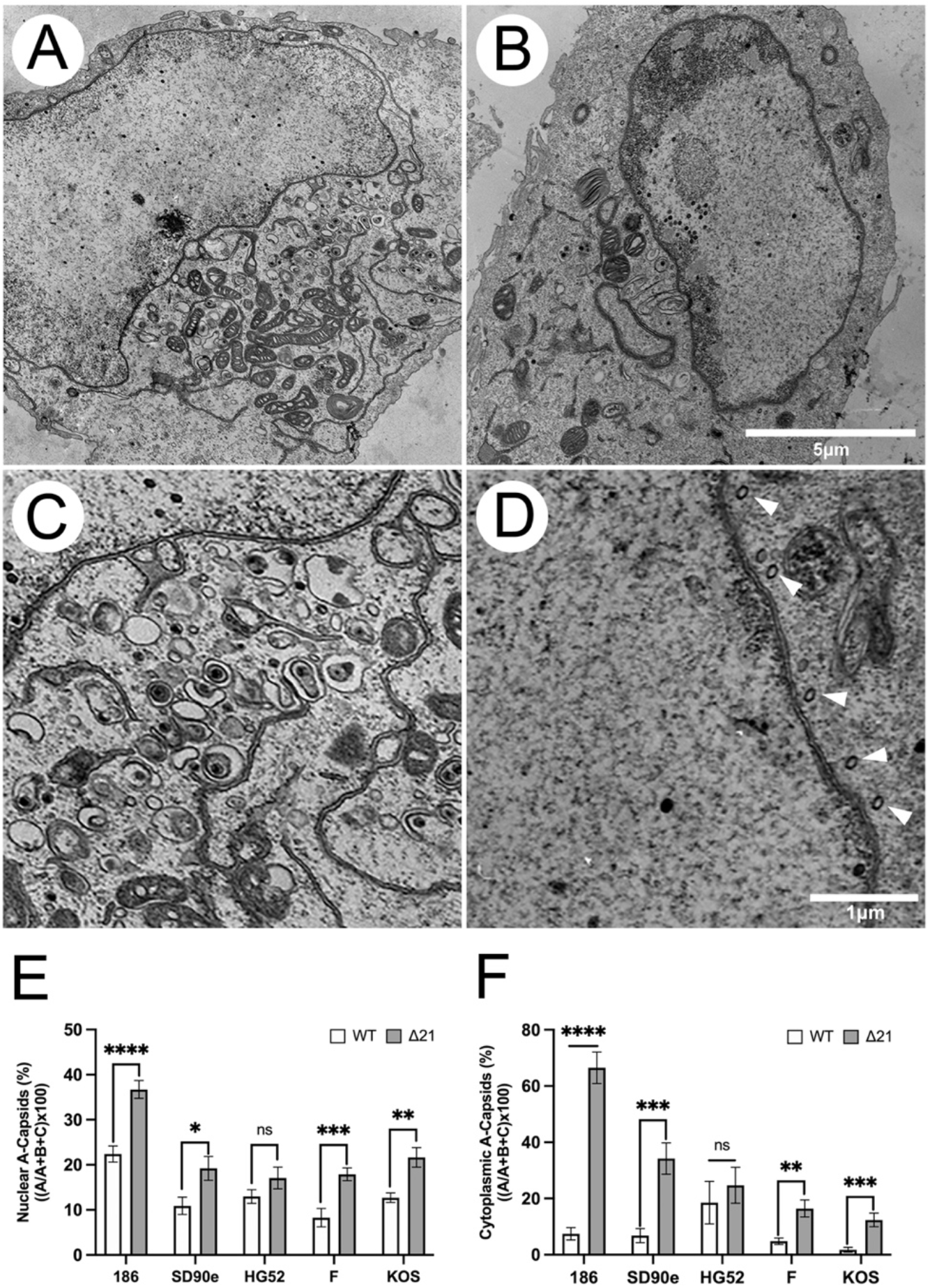
Nuclear and cytoplasmic A-capsid accumulation in HSV WT and Δ21 strains. Vero cells were infected with either an HSV-1 or HSV-2 WT virus or its corresponding Δ21 mutant. At 18 hpi, samples were fixed and prepared for TEM. Micrographs for each strain of HSV-1 and HSV-2 were obtained and WT and Δ21 capsids in the nucleus and cytoplasm of infected cells were quantified. **A-D)** Example micrographs of HSV-2 SD90e WT (**A** and **C**) compared to its Δ21 mutant (**B** and **D**). As depicted in the Δ21 panels (**B** and **D**), there is an abundance of A-capsids (white arrowheads) compared to the WT strain (**A** and **C**). The 5µm scale bar applies to the top two panels and the 1µm scale bar applies to the bottom two panels of the figure. **E)** The quantification of nuclear A-capsids in cells infected with HSV WT and Δ21 strains are depicted (n=9-12 micrographs per condition). **F)** The quantification of cytoplasmic A-capsids of cells were infected with HSV WT and Δ21 strains are depicted (n=11-14 micrographs per condition). Unpaired t-tests were conducted between Δ21 mutants and WT viruses. *, **, ***, and **** represent p ≤ 0.05, p ≤ 0.01, p ≤ 0.001 and p ≤ 0.0001, respectively. ns = not significant.

TEM micrographs were also used to quantify the number of A-capsids in the cytoplasm of infected cells. The percentage of cytoplasmic A-capsids in cells infected with HSV-2 (186, SD90e, HG52) and HSV-1 (F, KOS) WT viruses, along with their corresponding Δ21 mutants, were determined (Fig 1F). The Δ21 mutants of HSV-2 186 and SD90e, as well as HSV-1 F and KOS, had significantly more A-capsids present in the cytoplasm compared to the WT infections. There was no significant difference (p = 0.2701) between the HSV-2 HG52 Δ21 mutant and WT virus, despite there being more cytoplasmic A-capsids in Δ21 infected cells.

These data demonstrate that cells infected with Δ21 mutants from all HSV-1 strains tested and two out of three HSV-2 strains have significantly more nuclear and cytoplasmic A-capsids compared to WT infected cells. Considering more cytoplasmic A-capsids are present during Δ21 infection (Fig 1F), we next determined if Δ21 extracellular virions contained more A-capsids than WT extracellular virions.

### Deletion of UL21 decreases the proportion of genome-containing capsids within extracellular virions

Previous studies examining PRV mutants that cause rupture of the nuclear envelope and leakage of A-, B-, and C-capsids into the cytoplasm found that these mutants efficiently incorporated A- and B-capsids into extracellular virions (42, 43). This suggests that during PRV infection, C-capsids are not preferentially selected for secondary envelopment. To assess if this was also true for HSV-1 and HSV-2, we quantified the proportion of WT and Δ21 extracellular virions containing viral genomes.

Recombinant WT and Δ21 strains expressing red fluorescent proteins fused to the capsid protein VP26 were used to determine if HSV capsids lacking viral genomes were packaged into extracellular virions and how the absence of pUL21 affected the proportion of empty capsids within these virions. Extracellular virions were stained with the green-fluorescent nucleic acid dye, syto13, and examined by confocal microscopy. Empty capsids (A- or B-capsids) were identified by a red signal alone, while genome-containing capsids (C-capsids) were identified by a yellow signal indicating the colocalization between viral genome (green) and capsid (red) signals (Fig 2A-D). HSV-1 and HSV-2 Δ21 mutants had significantly more empty capsids in extracellular virions compared to corresponding WT viruses (Fig 2E). The data suggest that WT and Δ21 cytoplasmic empty capsids (A- and/or B-capsids) are readily enveloped and that more empty capsids are found in extracellular virions in the absence of pUL21.

**Fig 2.**
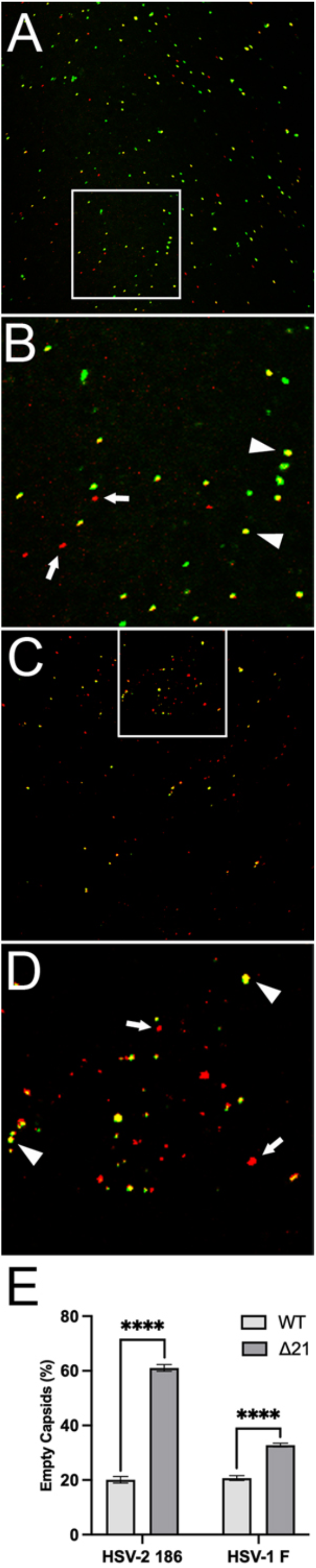
Comparison of extracellular virions containing empty capsids released from HSV WT and Δ21 infected cells. **A-D)** Viral genomic DNA inside WT HSV-2 186 mCh labelled capsids (**A** and **B**) or 186 Δ21 mChVP26 labelled capsids (**C** and **D**) were stained with the green-fluorescent dye, syto13. Capsids were analyzed by confocal microscopy to identify empty capsids (A- and B-capsids) present within each virion population. Capsids were counted using Image Pro Plus version 6.3.0.512. The yellow signal represents C-capsids (arrowheads), the red signal represents empty capsids (arrows), and the green signal represents viral or cellular DNA attached to the poly-L-lysine coated glass-bottom dish. C-capsids appear yellow due to the co-localization of red-capsid and DNA signals. **E)** Quantification of HSV-2 186 and HSV-1 F empty capsids within extracellular virions is depicted (n=30 images per condition). Unpaired t-tests were conducted between Δ21 mutant and WT viruses for three biological replicates. **** represents p ≤ 0.0001.

### Nuclear integrity remains intact in Δ21 infected cells

The data described thus far demonstrated that, in the absence of pUL21, there are more nuclear, cytoplasmic, and extracellular A-capsids, suggesting that genome retention within capsids was impaired. However, if nuclear envelope integrity was compromised during Δ21 infection, passive leakage of A-capsids from the nucleoplasm into the cytoplasm may occur. To evaluate nuclear integrity during Δ21 infection, Vero cells were transfected with pRF52, a plasmid encoding EGFP-JUMBO, a 133 kDa fusion protein that localizes to the cytoplasm, lacks a nuclear import signal, and is too large to passively diffuse through nuclear pore complexes. Thus, the subcellular localization of this protein can be used to assess nuclear envelope integrity. Twenty-four hours post-transfection with pRF52, cells were mock-infected or infected with HSV-1 F WT or Δ21 for 24 hours. EGFP signal was mostly cytoplasmic in mock, WT and Δ21 infected cells (Fig 3A) and quantification of nuclear:cytoplasmic ratios of EGFP-JUMBO fluorescence intensity revealed no significant difference (p = 0.6121) between HSV-1 F WT and Δ21 infected cells (Fig 3B). These data indicate that the integrity of the nuclear envelope remains intact in WT and Δ21 infected cells and strongly suggest that A-capsid accumulation in the cytoplasm of Δ21 infected cells was not due to the loss of nuclear envelope integrity during infection. In further support of this conclusion, we failed to identify B-capsid accumulation in the cytoplasm of Δ21 infected cells (Fig 1), arguing against gross perturbations of nuclear envelope barrier function in these cells.

**Fig 3.**
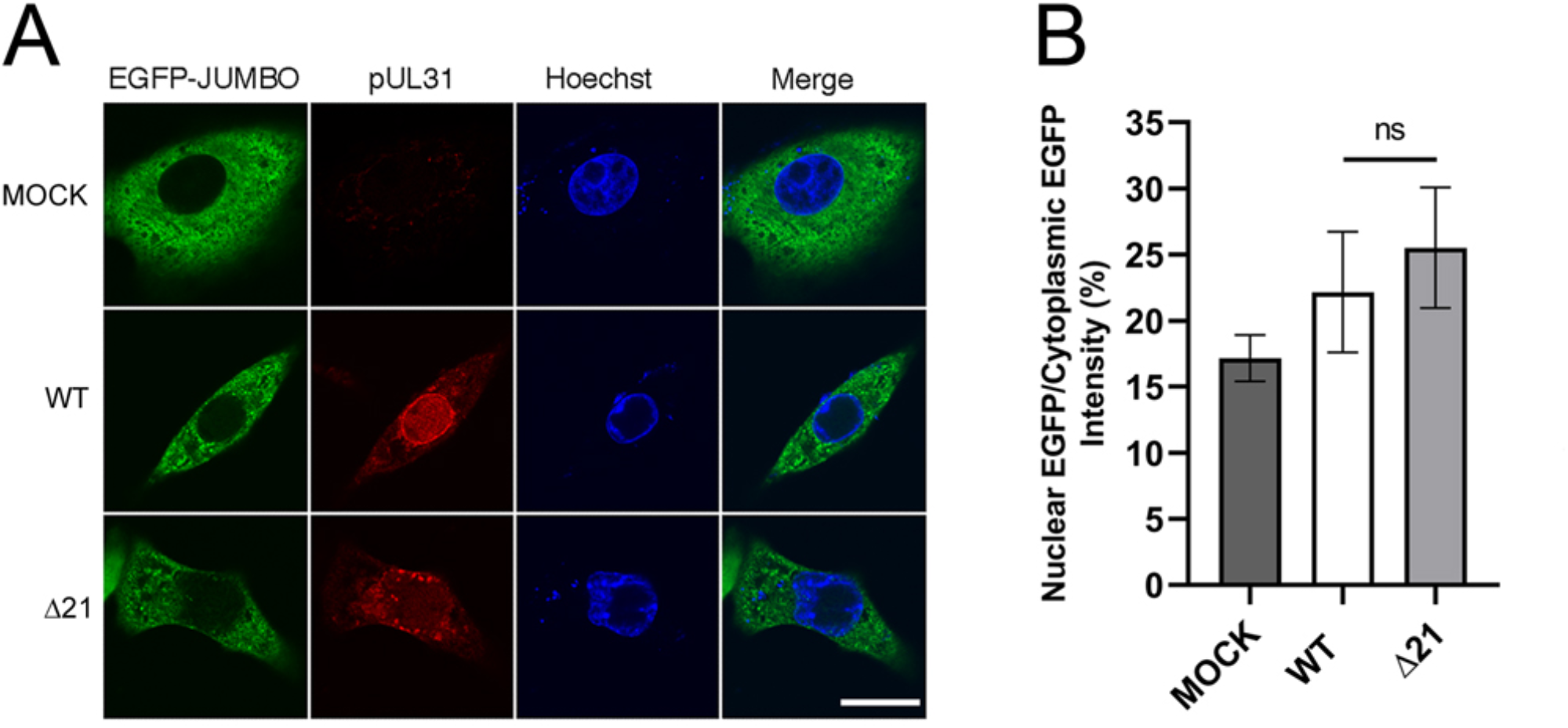
Nuclear integrity remains intact in ΔUL21 infected cells. **A)** Confocal images of cells transfected with pRF52, which directs the production of EGFP-JUMBO, a 133 kDa EGFP fusion protein that has a predominantly cytoplasmic localization. At 24 hours post-transfection cells were mock-infected or infected with HSV-1 F WT or HSV-1 F Δ21, for 24 hours prior to fixing and staining cells for pUL31 (red) to identify infected cells by confocal microscopy. The scale shown is 20μm and applies to all panels. **B)** Quantification of nuclear:cytoplasmic ratios of EGFP-JUMBO fluorescence intensity for n = 7 cells per condition. Pixel intensities were measured in the nuclei and cytoplasm of transfected cells using FluoView software version 4.01. The differences in nuclear:cytoplasmic ratios of EGFP-JUMBO were not significant between cells infected with WT and Δ21 strains (p = 0.6121).

### Preferential selection of C-capsids for nuclear egress is operational in the absence of pUL21

Another possible explanation for the increased numbers of A-capsids in the cytoplasm of Δ21 infected cells is that there is a breakdown in C-capsid selectivity during nuclear egress such that A-capsids acquire primary envelopes efficiently, enabling their delivery to the cytoplasm. To address this, we quantified primary enveloped virions (PEVs) by TEM (Fig 4A). If C-capsid selectivity was functional in the absence of pUL21, C-capsids should be preferentially found within PEVs. Capsids inside PEVs were scored as A-capsids, B-capsids, C-capsids, or unidentified (i.e., capsids that could not be unambiguously identified). Minimal differences were seen in the percentage of C-capsids within PEVs when comparing HSV-1 (F, KOS) and HSV-2 (186, SD90e, HG52) Δ21 mutants to their corresponding WT viruses (Fig 4B). Indeed, most Δ21 strains had more C-capsids within PEVs. This may be due to the accumulation of PEVs in the perinuclear space, previously observed in the absence of pUL21 (27). Nevertheless, these analyses demonstrated that the selectivity of C-capsids for nuclear egress remained functional in the absence of pUL21 and that enhanced selection of A-capsids for primary envelopment cannot explain the accumulation of A-capsids in the cytoplasm.

**Fig 4.**
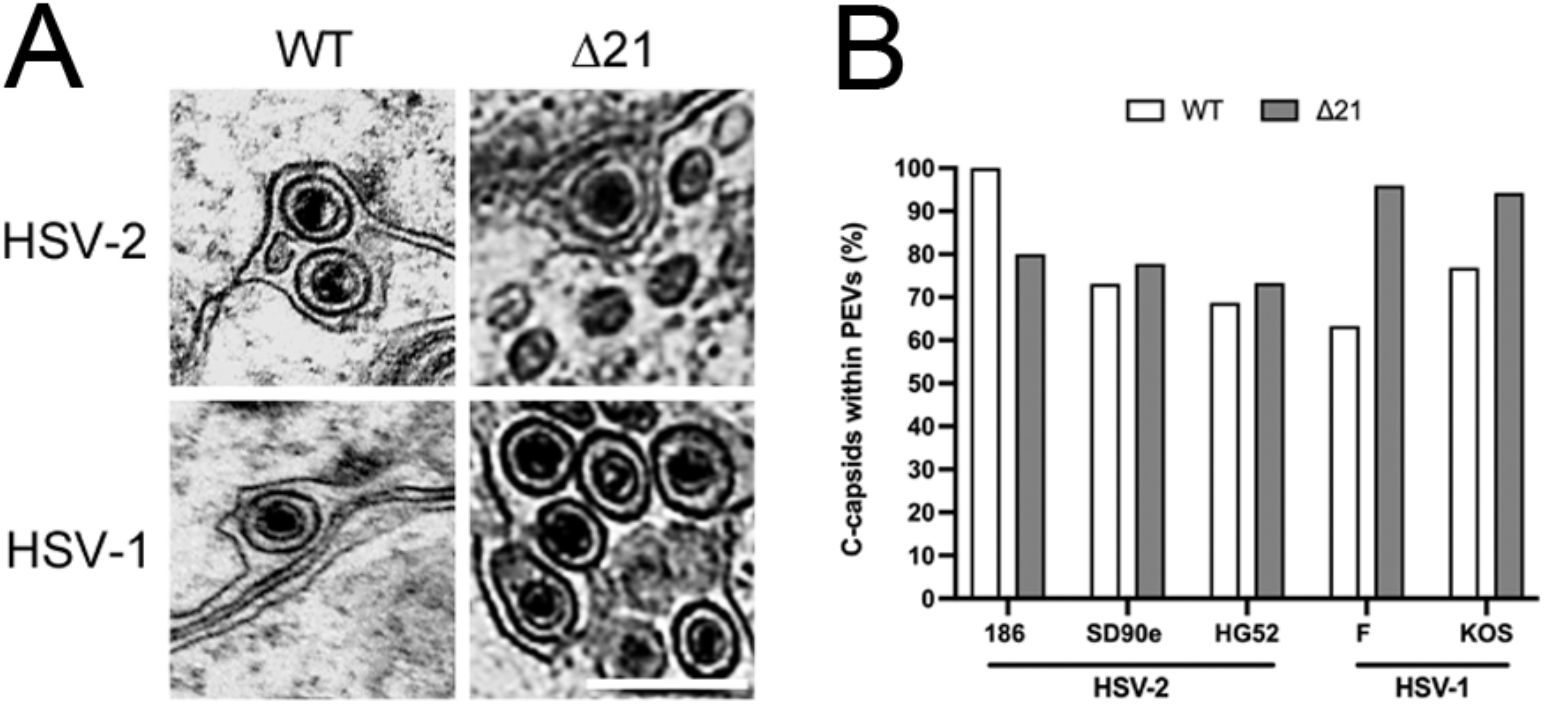
Perinuclear C-capsid abundance in HSV WT and Δ21 infected cells. Vero cells were infected with HSV-1 or HSV-2 WT strains or their corresponding Δ21 mutants for 18 h. Cells were then fixed and prepared for TEM. Micrographs for each strain of HSV-1 and HSV-2 were obtained to quantify WT and Δ21 capsids within PEVs. **A)** Examples of micrographs containing PEVs derived from HSV-2 SD90e WT and its Δ21 mutant and HSV-1 F WT and its Δ21 mutant. The scale bar shown is 500nm and applies to all panels. **B)** The quantification of C-capsids within PEVs from cells infected with HSV WT and Δ21 strains (n = 11-27 micrographs per condition). Statistical analysis was not possible due to PEVs being a transient assembly intermediate and not consistently present within each micrograph.

### Viral genome retention within capsids is compromised in the absence of pUL21

To evaluate WT and Δ21 genome retention within capsids, we examined genome ejection during the early stages of infection using click-chemistry. Early in infection, after fusion of the virion envelope with a cellular membrane, capsids containing viral genomes are transported to nuclear pore complexes through which genomes are deposited into the nucleus. However, if capsids fail to retain their genomes, they are prematurely ejected into the cytoplasm, which can be quantified. HSV-1 KOS WT and Δ21 strains were propagated in the presence of the nucleoside analog, EdC, which was incorporated into the genomes of progeny virions. Viruses containing EdC-labelled genomes were then used to infect Vero cells growing on glass-bottom dishes. Cells were fixed at intervals between 0 and 4 hpi and click-chemistry was performed to fluorescently label EdC within viral genomes that had been ejected from capsids (Fig 5A). Importantly, EdC-labelled genomes that were contained within capsids were not fluorescently labelled by the click reaction (Fig 5B). Fluorescent puncta, representing individual viral genomes, were quantified to determine the number of nuclear (Fig 5C) and cytoplasmic (Fig 5D) viral genomes released from capsids per cell, over time. Δ21 infected cells had significantly fewer viral genomes delivered to nuclei between 1 and 4 hpi compared to WT (Fig 5C). Additionally, Δ21 infected cells had significantly more non-encapsidated viral genomes in the cytoplasm between 1 and 4 hpi compared to WT (Fig 5D). These findings indicate that HSV-1 KOS Δ21 capsids have impaired viral genome retention, explaining the abundance of A-capsids in Δ21 infected cells.

**Fig 5.**
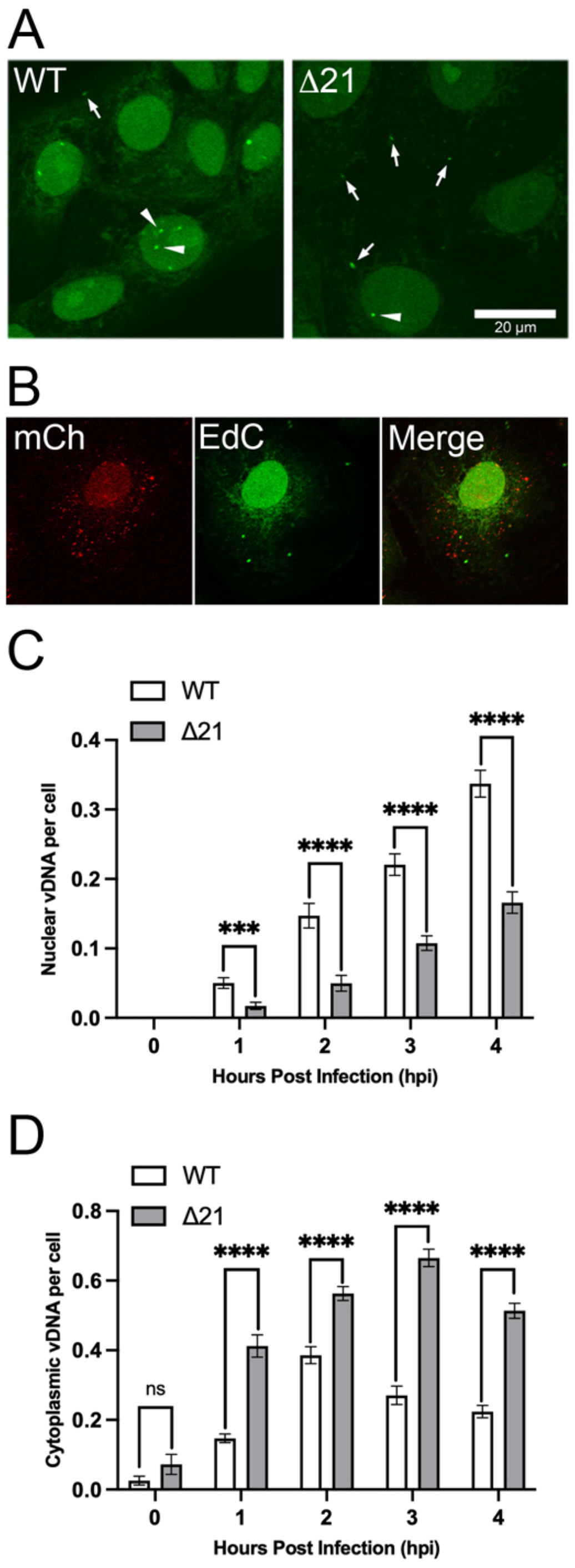
Comparison of genome retention abilities of HSV-1 KOS WT and Δ21 capsids during early infection. Vero cells were infected at an MOI of 0.1 with HSV-1 KOS WT or Δ21 viruses containing EdC incorporated into their genomes. Infected cells were fixed at early times after infection (0-4 hpi) and genomes ejected from capsids were labeled using click-chemistry. **A)** Representative images of WT and Δ21 infected cells at 4 hpi with ejected viral genomes labeled using click-chemistry. The scale bar is 20μm. Labelled genomes were identified as intense puncta seen in the cytoplasm (arrows) and nuclei of infected cells (arrowheads). Puncta were quantified and their numbers divided by the total number of cells to determine nuclear and cytoplasmic genomes per cell. **B)** Control experiment using EdC labeled HSV-2 that produces mCh labeled nucleocapsids showing that click-chemistry does not label genomes within capsids. **C)** The quantification of nuclear viral genomes (vDNA) per cell over 0-4 hpi is shown (n=21 images per timepoint and condition). **D)** The quantification of cytoplasmic genomes (vDNA) per cell over 0-4 hpi is shown (n=21 images per timepoint and condition). Unpaired t-tests were performed between Δ21 mutants and WT viruses for three biological replicates. *** and **** represent p ≤ 0.001 and p ≤ 0.0001, respectively.

As a complementary approach to analyze premature ejection of viral genomes in other HSV-1 and HSV-2 Δ21 mutants, activation of the cGAS/STING pathway was examined. When viral genomes are prematurely ejected into the cytoplasm, they can be detected by components of the cGAS/STING pathway leading to interferon regulatory factor 3 (IRF-3) phosphorylation in the cytoplasm and its subsequent translocation to the nucleus (44, 45). Thus, nuclear translocation of IRF-3 can be used as a surrogate measure of premature ejection of viral genomes into the cytoplasm of infected cells (45, 46). Due to the severe growth defect of HSV-2 186 Δ21 virus (41), this strain was not investigated in these experiments. T12 cells, life extended human fibroblasts, were used for these experiments because, unlike Vero cells, they can produce interferon and thus have intact IRF-3 signalling pathways. T12 cells were infected with WT or Δ21 viruses at an MOI of 3. Cells with nuclear IRF-3 were quantified in WT and Δ21 infected cells by confocal microscopy (Fig 6A). At 2, 4, and 6 hpi, HSV-1 F and KOS Δ21 infections had significantly more cells with nuclear IRF-3 compared to their respective WT infections (Figs 6B, 6C). Additionally, during HSV-1 F and KOS Δ21 infections, translocation of IRF-3 to infected nuclei occurred 2 hours earlier than their respective WT infections. At 2, 4, and 6 hpi, HSV-2 SD90e Δ21 infection also displayed significantly more cells with nuclear IRF-3 compared to WT (Fig 6D). Collectively, these data suggest that Δ21 mutants prematurely eject more viral genomes into the cytoplasm (Figs 5D, 6A-D) and deliver fewer genomes to the nuclei of infected cells (Fig 5C).

**Fig 6.**
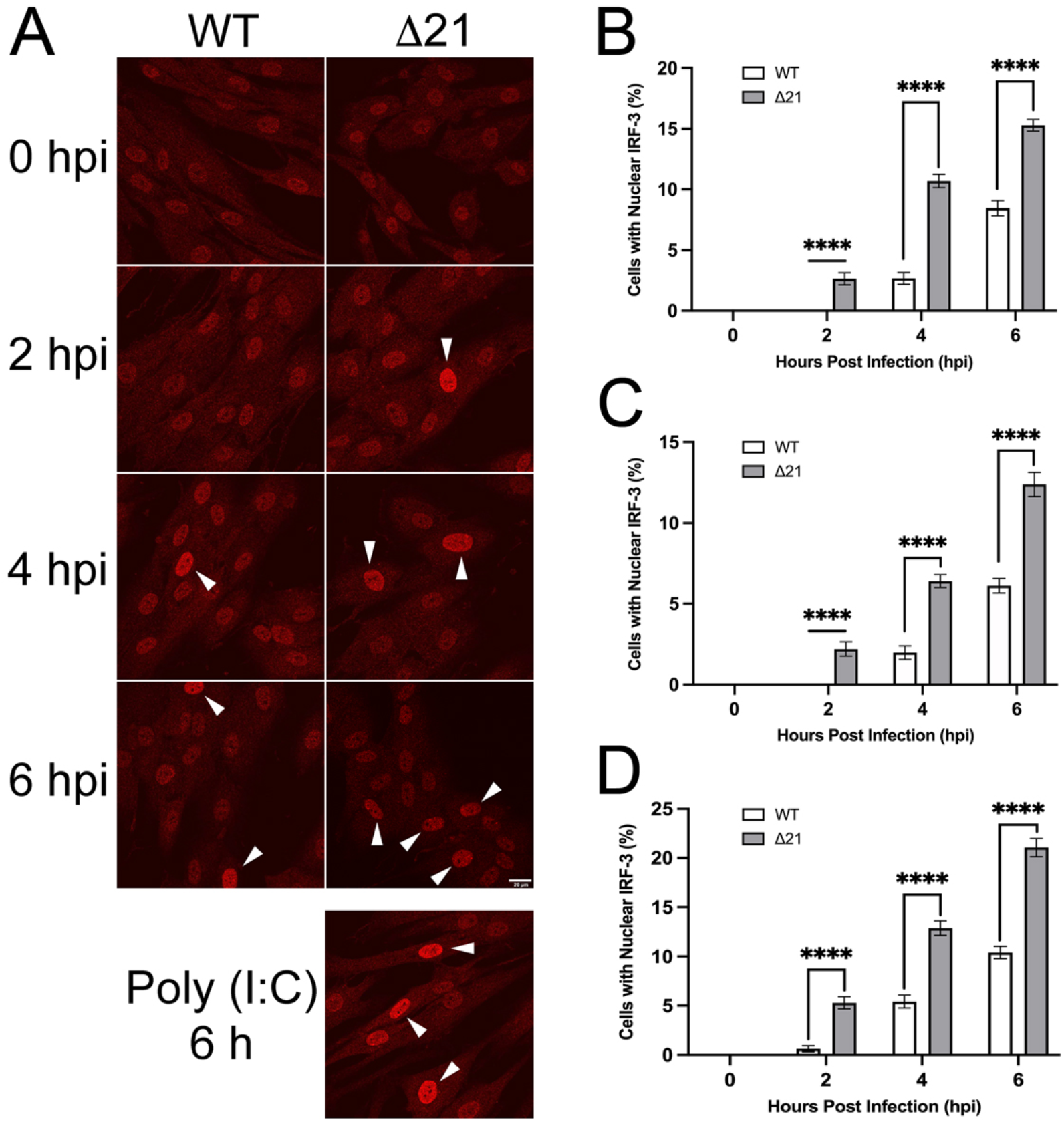
Examination of IRF-3 nuclear localization during early stages of infection with HSV WT and Δ21 strains. T12 cells were infected at an MOI of 3 and after inoculation for one-hour on ice, cells were incubated in medium containing 50 μg/ml of cycloheximide to inhibit protein synthesis. Infected cells were fixed at 0, 2, 4 and 6 hpi, stained for IRF-3 and imaged by confocal microscopy. **A)** Representative images of HSV-1 KOS WT and Δ21 infected cells. IRF-3 localization at 0, 2, 4, and 6 hpi. Nuclear IRF-3 is indicated by the white arrowheads. Representative image of T12 cells transfected for 6 hours with poly I:C is shown as a positive control. n = 33-37 images per timepoint and condition. Scale bar is 20μm and applies to all panels. **B)** Quantification of nuclear IRF-3 in cells infected with HSV-1 KOS WT and Δ21 strains (n = 33-36 images per timepoint and condition). **C)** Quantification of nuclear IRF-3 in cells infected with HSV-1 F WT and Δ21 strains (n=36 images per timepoint and condition). **D)** Quantification of nuclear IRF-3 in cells infected with HSV-2 SD90e WT and Δ21 strains (n=36 images per timepoint and condition). Unpaired T-tests were performed to compare the percentage of cells with nuclear IRF-3 for each timepoint in Δ21 mutants with WT virus for each strain, from three biological replicates. **** represents p ≤ 0.0001.

### The composition of nuclear capsids is altered in the absence of pUL21

Previous studies examining viruses mutated for capsid and tegument proteins have reported changes in capsid and tegument composition (7, 14, 39, 47-49). To understand the mechanism by which Δ21 capsids have impaired genome retention we investigated capsid composition by mass spectrometry. After the exclusion of common contaminants, a total of 265 unique cellular and viral proteins were identified in HSV-1 KOS WT and Δ21 nuclear A-, B-, and C-capsid preparations (Table S1). Variations between capsid abundance in different biological replicates as well as between WT and Δ21 strains were normalized using the total spectrum count abundance of the major capsid protein, VP5, which is found in exactly 955 copies per capsid (50). Next, the percent change of each identified protein was determined and averaged over the three biological replicates for Δ21 A-, B-, and C-capsids compared to WT A-, B-, and C-capsids. We defined a significant protein change as being greater than a 20% increase, or decrease, in spectrum count abundance. Of the 265 identified proteins, the tegument protein pUL16 was increased by 9.68%, 96.52%, and 161.75% on Δ21 A-, B-, and C-capsids, respectively, compared to WT capsids. pUL16 is a known binding partner of pUL21 and, interestingly, its association with capsids is thought to occur only in the cytoplasm of infected cells (32, 51). The CVSC is known to be vital for capsid stability and genome retention (14). The amount of the CVSC proteins, pUL17 and pUL25, were similar when comparing Δ21 capsids to WT capsids. Interestingly, the CVSC protein pUL36 was decreased by 37.19% on Δ21 A-capsids compared to WT A-capsids (Table S1). Unlike pUL25, pUL36 has not been implicated in genome retention and pUL36 mutants have a similar amount of nuclear A-, B-, and C-capsids compared to WT infected nuclei (14). Given this, it is unlikely that alterations in pUL36 levels are impairing genome retention within capsids in the absence of pUL21. Furthermore, no cellular proteins that met our defined criteria stood out as candidates for impairing capsid genome retention in the absence of pUL21. Thus, we next examined if pUL16 was increased on a variety of HSV-1 and HSV-2 Δ21 capsids by western blotting.

### Increased levels of pUL16 are associated with HSV-1 and HSV-2 nuclear capsids in the absence of pUL21

Nuclear capsids were isolated from infected cells by ultracentrifugation through 20-50% sucrose gradients to resolve A-, B-, and C-capsids (Fig 7A). No apparent differences were seen when comparing the light-scattering bands of Δ21 A-, B-, and C-capsids to the light-scattering bands of WT A-, B-, and C-capsids. Sucrose gradients were fractionated, and capsid-containing fractions were subjected to SDS-PAGE followed by silver-staining (Fig 7B). No obvious differences in protein profiles were seen between WT and Δ21 capsids. WT and Δ21 capsids were then analyzed by western blotting using antisera reactive against VP5, pUL17, pUL25, pUL21, and pUL16. Blots were imaged using a LI-COR CLx imager to quantify differences in capsid protein composition.

**Fig 7.**
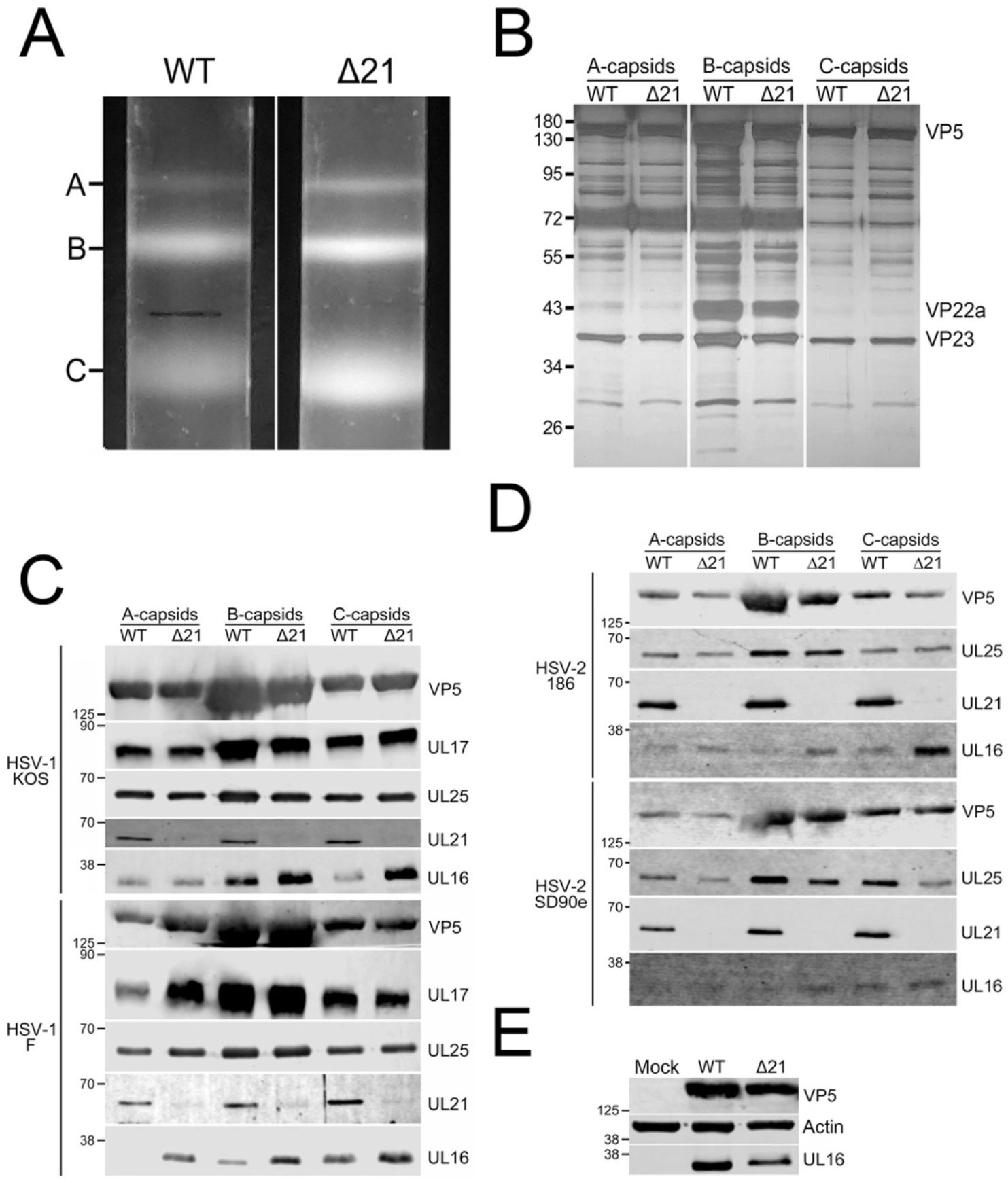
Isolation and analysis of HSV WT and Δ21 nuclear A-, B-, and C-capsids. Nuclear capsids were isolated from Vero cells infected with HSV-1 or HSV-2 WT and corresponding Δ21 strains. **A)** Light scattering bands observed in 20-50% sucrose gradients used to resolve A-, B- and C-capsids isolated from cells infected with WT and Δ21 HSV-1 KOS. **B)** Nuclear capsid-containing gradient fractions from WT and Δ21 HSV-1 KOS infected cells were electrophoresed through an SDS-PAGE gel and silver-stained to identify proteins associated with A-, B-, and C-capsids. VP5 and VP23 can be clearly seen in all samples, while VP22a is only seen in WT and Δ21 B-capsid fraction samples. No obvious differences were observed between WT and Δ21 A-, B-, and C-capsids **C and D)** Western blot analysis of HSV-1 (KOS, F) and HSV-2 (186, SD90e) WT and Δ21 A-, B-, and C-capsid fractions probed with antisera, indicated on the right side of each panel. Migration positions of molecular weight markers (kDa) are indicated on the left side of each panel. **E)** Vero cells were mock-infected or infected with HSV-1 KOS WT or Δ21 strains. At 18 hpi, cell lysates were analyzed by western blotting using the antisera indicated on the right side of each panel. The migration position of molecular weight markers (kDa) are indicated on the left side of each panel.

Capsid loading levels were normalized using the signal obtained from the major capsid protein VP5. While there were modest fluctuations in the amounts of pUL17 and pUL25 present in A-, B- and C-capsids from WT and Δ21 HSV-1 and HSV-2 strains, none of these fluctuations were consistent between strains (Figs 7C, 7D, Table 1). For example, while A-capsids from HSV-1 F Δ21 had more pUL17 than the parental WT strain, this was not the case for HSV-1 KOS (Fig 7C). Likewise, while C-capsids from HSV-2 SD90e Δ21 had less pUL25 than the parental WT strain (Fig 7D), this was not the case for the other HSV-2 and HSV-1 strains investigated. The only difference in capsid composition consistently observed between all HSV-1 and HSV-2 strains analyzed was an increase in pUL16 associated with Δ21 capsids in comparison to their WT counterparts.

**Table 1.** Quantitative analysis of capsid associated proteins.

| Strain | Protein | Number of Biological Replicates | Average Protein Change (%) ($\Delta 21$ compared to WT)<sup>a</sup> | | |
| --- | --- | --- | --- | --- | --- |
|  |  |  | A-capsid | B-capsid | C-capsid |
| HSV-1 KOS | pUL17 | 3 | -18.85 ± 29.97 | 1.08 ± 6.41 | 0.28 ± 20.68 |
|  | pUL25 |  | -5.97 ± 29.20 | 6.86 ± 19.00 | -2.28 ± 9.36 |
|  | pUL16 |  | 79.99 ± 33.48 | 169.22 ± 89.55 | 197.22 ± 150.10 |
| HSV-1 F | pUL17 | 2 | 85.87 ± 11.08 | 3.97 ± 3.00 | 7.71 ± 17.23 |
|  | pUL25 |  | 16.80 ± 17.09 | -11.64 ± 7.87 | 38.81 ± 5.46 |
|  | pUL16 |  | 189.63 ± 8.15* | 133.83 ± 13.28* | 83.07 ± 26.41 |
| HSV-2 186 | pUL25 | 1 | 4.71 | 12.26 | 40.25 |
|  | pUL16 |  | 108.57 | 274.78 | 663.05 |
| HSV-2 SD90e | pUL25 | 1 | -12.41 | -36.55 | -30.37 |
|  | pUL16 |  | 156.69 | 26.46 | 28.83 |

<sup>a</sup> The mean ± standard error of the mean is shown. For samples with only one biological replicate, only the mean is shown.
\* p<0.05

A trivial explanation for these findings is that increased pUL16 expression in the absence of pUL21 led to enhanced pUL16 association with capsids. To eliminate this possibility, we examined pUL16 expression levels in Δ21 infected cells (Fig 7E). WT and Δ21 infected cell lysates were normalized using VP5 signal intensity. Δ21 infected cells had 75% less pUL16 compared to WT infected cells. This suggests that the increased association of pUL16 with Δ21 nuclear capsids was not due to elevated expression levels of pUL16 in Δ21 infected cells.

### Deletion of UL16 from HSV-1 KOS Δ21 decreased the abundance of nuclear A-capsids in infected cells

We reasoned that if the increased association of pUL16 with Δ21 nuclear capsids was impairing genome retention, then deletion of UL16 from HSV-1 KOS Δ21 strain would restore genome retention. We used CRISPR/Cas9 mutagenesis to construct two independent HSV-1 KOS Δ21/Δ16 mutant strains (HΔ21/Δ16 and EΔ21/Δ16). These strains were analyzed by TEM and numbers of A-, B- and C-capsids were quantified in the nuclei of infected cells (Fig 8A-D). Interestingly, limited numbers of cytoplasmic capsids were seen in Δ21/Δ16 infected cells, suggesting that the combined effect of deleting both UL21 and UL16 genes led to decreased nuclear egress. Whilst the Δ21 mutant had roughly twice as many A-capsids in the nucleus as the parental WT strain, both Δ21/Δ16 double mutants had numbers of A-capsids that were indistinguishable from the WT strain. These findings demonstrate that the absence of both pUL16 and pUL21 restored the ability of capsids to retain genomes at WT levels. These data, coupled with our findings that HSV-1 and HSV-2 Δ21 mutants have more capsid-associated pUL16 compared to WT capsids, suggest that increased and/or premature association of pUL16 with nuclear capsids in the absence of pUL21 impairs capsid genome retention.

**Fig 8.**
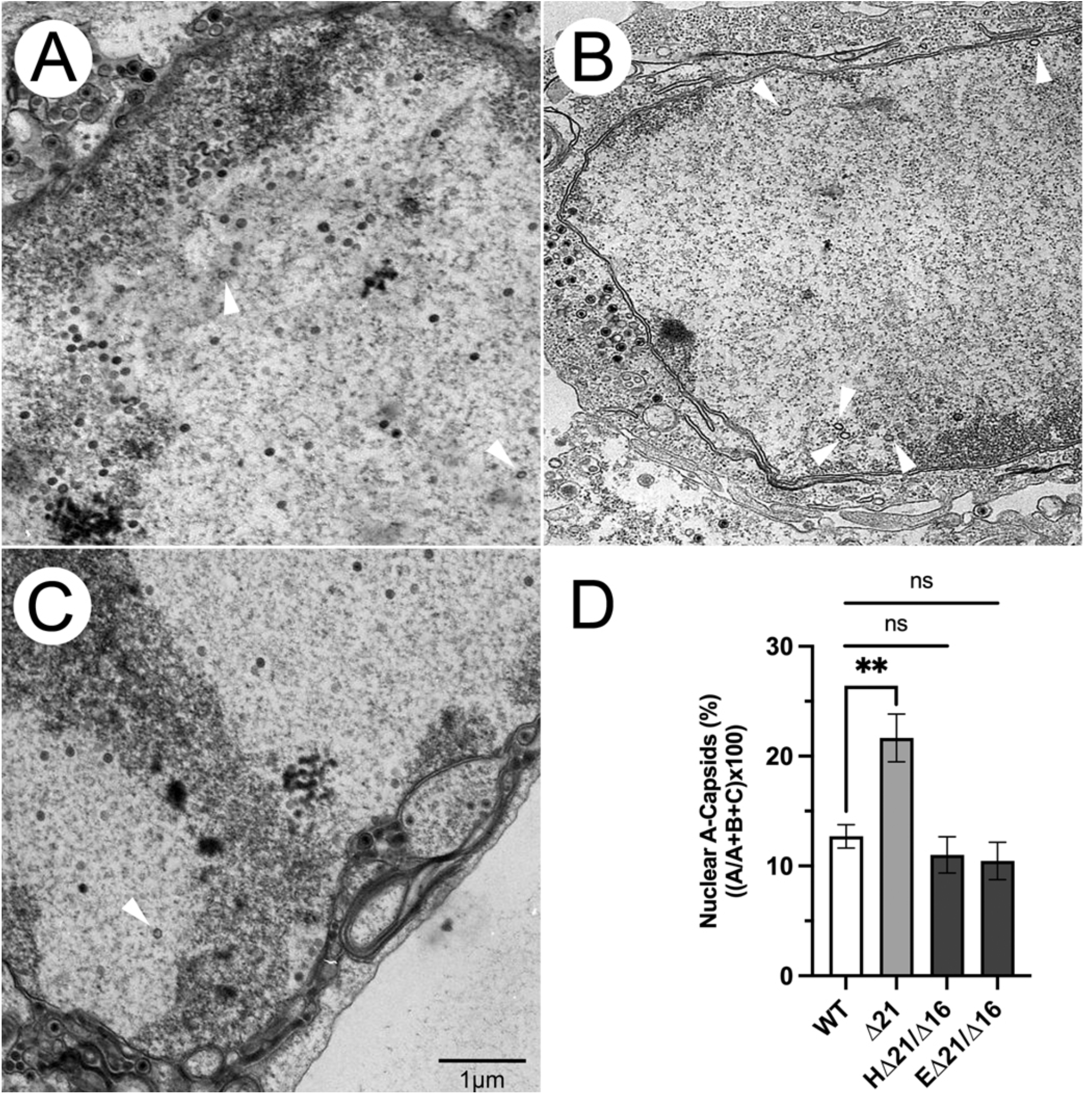
Characterization of capsids produced by Δ21/Δ16 double mutants. **A-C)** Vero cells were infected with either HSV-1 KOS WT (**A**), Δ21 (**B**), or EΔ21/Δ16 strains (**C**) for 18 hpi and examined by TEM. **D)** Quantification of nuclear A-capsids in cells infected with the indicated HSV-1 KOS derived strains. n = 11-14 micrographs per condition. Unpaired t-tests were performed between Δ21 mutant and WT, Δ21/Δ16 and WT, as well as Δ21/Δ16 and Δ21 strains. ** p ≤ 0.01. The difference in nuclear A-capsids was not significant (p = 0.2078) when comparing Δ21/Δ16 mutants to WT.

## Discussion

In this study we have defined the mechanism by which A-capsids accumulate in cells infected with multiple HSV-1 and HSV-2 UL21 deletion mutants. We found that A-capsids accumulated in the cytoplasm and nuclei of Δ21 infected cells (Fig 1) and that capsids lacking genomes were more commonly incorporated into Δ21 extracellular virions compared to WT viruses (Fig 2). Interestingly, despite HSV-2 strain HG52 having more nuclear and cytoplasmic A-capsids present in Δ21 infected cells compared to WT infected cells, this was the only strain where the differences between WT and Δ21 were not significant (Fig 1). Given that the pUL21 proteins from HSV-2 186, SD90e and HG52 are identical, it was surprising that strain HG52 was an outlier. When comparing the proportion of nuclear and cytoplasmic A-capsids from each WT infection, HG52 infected cells had the second highest percentage of nuclear A-capsids (Fig 1B) and the highest percentage of cytoplasmic A-capsids (Fig 1C).This observation suggests that genome packaging, or the formation of HG52 capsids, is inherently defective relative to the other strains investigated and that the absence of pUL21 might not have as great an impact on A-capsid accumulation as it would in other strains. It may be that HG52 has polymorphisms outside the UL21 locus that influence capsid formation or genome retention/packaging. It is perhaps noteworthy that while pUL16 is identical in HSV-2 strains 186 and SD90e there is a stretch of five amino acids between residues 182 and 188 that differ in HG52 pUL16. It may be that HG52 pUL16 has a propensity to prematurely associate with nuclear capsids in the presence of pUL21; however, this needs to be addressed experimentally.

Our experiments indicated that A-capsid accumulation in the cytoplasm of Δ21 infected cells was not due to nuclear envelope rupture (Fig 3) or a breakdown in the preferential selection of C-capsids for nuclear egress (Fig 4). Rather, the data suggest that the ability of capsids to retain viral genomes is compromised in the absence of pUL21 (Figs 5, 6). These findings prompted an analysis of Δ21 capsid composition. Curiously, when nuclear A-, B-, and C-capsids were isolated by ultracentrifugation through sucrose gradients, WT A-capsid bands had a similar intensity to those derived from Δ21 infected nuclei (Fig 7A). Based on the increased proportion of A-capsids seen in the nuclei of Δ21 infected cells by TEM (Fig 1), we might have expected to see a more intense A-capsid band in the Δ21 samples. One explanation for this apparent discrepancy is that the bulk of the A-capsids seen in Δ21 infected cells by TEM are unstable and are not amenable to isolation on sucrose gradients. In support of this concept, cells infected with an HSV-1 pUL25 mutant produce nuclear A-, B-, and C-capsids when assessed by TEM, however, no C-capsid band is present on sucrose gradients of nuclear capsid preparations; presumably, because capsids lacking pUL25 are unable to withstand the capsid isolation procedure (7, 14). Despite this conundrum, our analyses identified more pUL16 on all capsid types isolated from HSV-1 and HSV-2 Δ21 in comparison to WT capsids (Fig 7, Table 1). In WT infected cells, pUL16 is not thought to associate with capsids until they reach the cytoplasm (51). As pUL16 has a pan-cellular distribution, this suggests that a mechanism exists to prevent the association of pUL16 with capsids in the nucleus. A simple explanation for these findings is that complex formation between pUL21 and pUL16 in the nucleus prevents pUL16 from associating with capsids (Fig 9).

**Fig 9.**
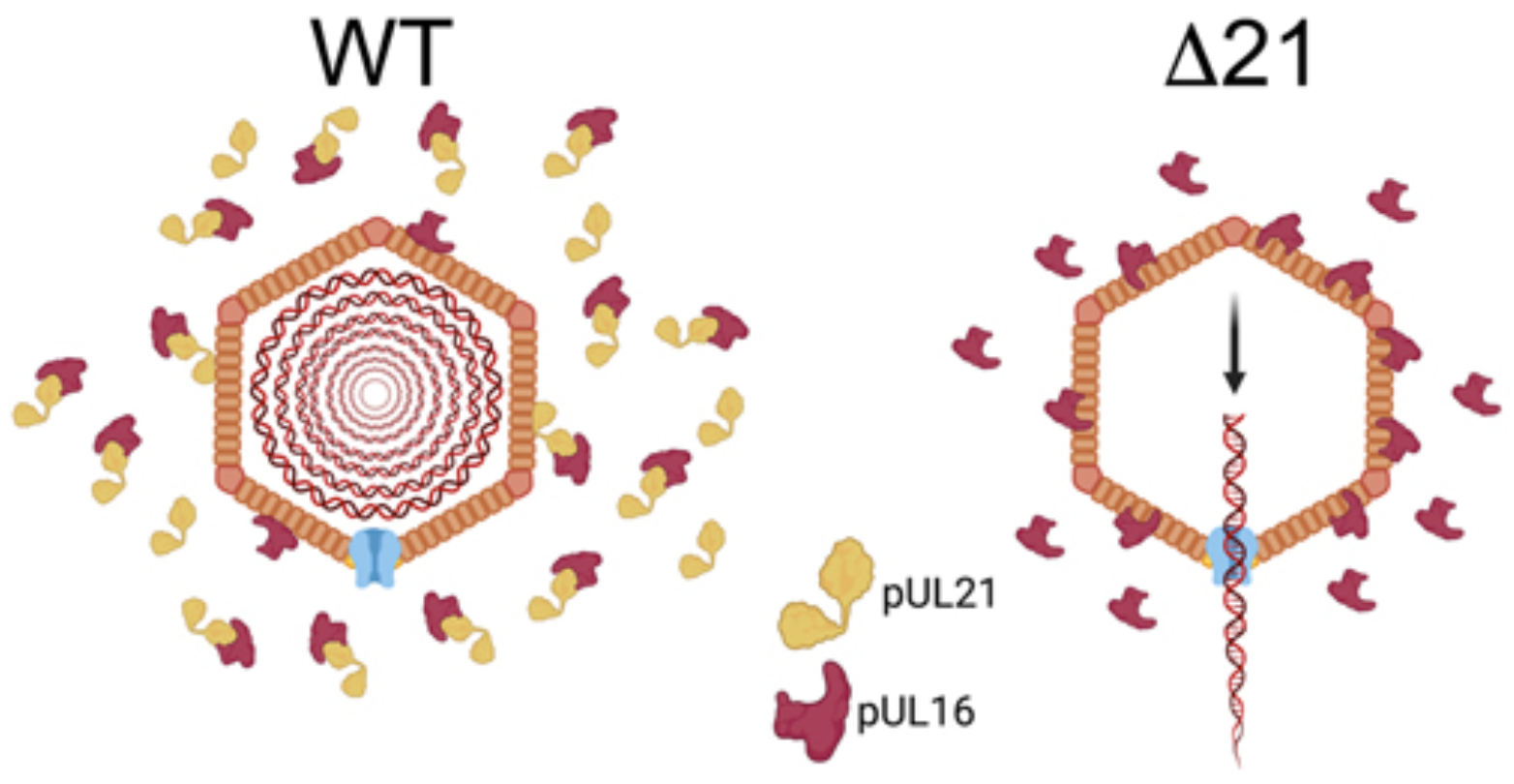
Possible mechanism accounting for impaired viral genome retention in the absence of pUL21. In the nuclei of cells infected with WT strains, sequestration of pUL16 by pUL21 prevents pUL16 from associating with nucleocapsids. However, in the absence of pUL21, pUL16 more readily associates with nucleocapsids and hinders genome retention within Δ21 capsids. The figure suggests that viral genomic DNA vacates Δ21 capsids via the portal, however, it may be that genomes escape these capsids at sites other than the portal. Figure created with BioRender.com.

To investigate this hypothesis, we constructed two independent HSV-1 KOS Δ21/Δ16 mutant strains (HΔ21/Δ16 and EΔ21/Δ16). We could only quantify nuclear A-capsids in HΔ21/Δ16 and EΔ21/Δ16 infected cells as there were limited numbers of cytoplasmic A-capsids in these cells. In HSV-1 mutants lacking UL21 or UL16 alone, nuclear egress defects have not been documented (37, 52-56). Thus, it was surprising that in HSV-1 KOS HΔ21/Δ16 and EΔ21/Δ16 infected cells, there were fewer cytoplasmic capsids than in HSV-1 Δ21 and WT infected cells, suggesting these double mutants have a nuclear egress defect. It may be that redundant activities within HSV-1 pUL21 and pUL16 can support nuclear egress whereas deletion of both molecules leads to deficiencies in this process. Interestingly, HSV-2 mutants deficient in pUL21, or pUL16 alone are deficient in nuclear egress (28, 57). Quantification of nuclear A-capsids in HΔ21/Δ16 and EΔ21/Δ16 infected cells resulted in no significant difference in the proportions of nuclear A-capsids compared to WT infection. However, there were significantly fewer nuclear A-capsids compared to Δ21 infected cells (Fig 8D). These findings indicate that the deletion of UL16 from the HSV-1 KOS Δ21 mutant restored capsid genome retention.

We hypothesize that pUL21 regulates the location and timing of pUL16 addition to capsids (Fig 9) and that during WT infection, pUL16 interaction with pUL21 in the nucleus prevents pUL16 from binding to capsids. How might premature and/or excessive pUL16 association with capsids impact capsid stability? During procapsid formation, the addition of capsomers and triplexes to the scaffolding proteins is regulated and occurs through interactions between pentons and hexons with scaffolding proteins and triplex components (16). Additionally, during capsid maturation, disulfide bond complexes form between capsid structural components (20-22). Specifically, disulfide bonds form between the major capsid protein VP5, triplex proteins, CVSC proteins, pUL6 and the tegument protein VP22. Disulfide bonds stabilize HSV capsids and disruption of these bonds can result in impaired capsid stability and genome retention. pUL16 contains 20 cysteines, with 8 of these conserved amongst all herpesviruses (51, 58). Thus, if pUL16 associates with nuclear capsids too early during assembly, it may result in improper disulfide bond formation between capsid components and impair genome retention after packaging. Future examination of disulfide bond complexes formed between capsid proteins in the absence of pUL21 should be conducted to better understand the mechanism(s) by which premature and/or over-addition of pUL16 affects genome retention within capsids.

In summary, this study has addressed a significant knowledge gap by elucidating the role of pUL21 in maintaining capsid stability in multiple HSV-1 and HSV-2 strains. The identification of factors that contribute to capsid stability exposes potential drug targets that can be exploited in the treatment of disease caused by these important human pathogens.

## Materials and Methods

### Cells and viruses

African green monkey kidney cells (Vero), life-extended human foreskin fibroblasts (T12), a kind gift from Dr. W. A. Bresnahan (University of Minnesota) (59), human keratinocytes (HaCaT), HaCaT21 and 293T21 cells, which stably express HSV-2 186 pUL21 (41), and HaCaT16 cells, which stably express HSV-2 186 pUL16 (27) were maintained in Dulbecco’s modified Eagle medium (DMEM) supplemented with 10% fetal bovine serum (FBS), penicillin/streptomycin and GlutaMax in a 5% CO_2_ environment. HSV-2 and HSV-1 strains deficient in pUL21 (Δ21) were constructed as described previously (28, 41). All pUL21 deficient viruses were propagated in HaCaT21 cells. Recombinant HSV-1 strain F containing monomeric red fluorescent protein (mRFP) fused to the minor capsid protein VP26 was a kind gift from Dr. G. A. Smith (Northwestern University). The corresponding Δ21 mutant was constructed by CRISPR-Cas9 mutagenesis as described previously (41). Recombinant HSV-2 strain 186 containing mCherry (mCh) fused to the minor capsid protein VP26 and the corresponding Δ21 mutant have been described previously (28). Times post-infection reported as hours post-infection (hpi) refers to the time elapsed following medium replacement after a one-hour inoculation period.

Two independently isolated HSV-1 KOS Δ21/Δ16 strains (HΔ21/Δ16 and EΔ21/Δ16) were constructed by CRISPR-Cas9 mutagenesis using UL16 guide RNAs reported previously (41). Briefly, HSV-1 KOS Δ21 genomic DNA was co-transfected into 293T21 cells along with plasmids expressing UL16-specific gRNAs using the calcium phosphate co-precipitation method (60). At 24 hours post-transfection, the medium was replaced with DMEM containing 2% FBS and 1% carboxymethylcellulose to enable plaque formation. Three days later, plaques were picked, and DNA was isolated from the picked plaques and screened for deletion of UL16 and UL21 by PCR. Viruses containing UL21 and UL16 deletions were analyzed by western blotting to confirm that pUL21 and pUL16 were not produced in infected cells. HSV-1 KOS HΔ21/Δ16 and EΔ21/Δ16 mutants were propagated on a 1:1 mixture of HaCaT21 and HaCaT16 cell monolayers.

### Plasmids

Plasmid pRF52 encodes a 133 kDa fusion protein, EGFP-JUMBO, comprised of WT HSV-2 Us3 fused to the N-terminus of a mutant form of human p53 that is defective in nuclear localization (R306A) (61) which in turn is fused to the N-terminus of EGFP.

### Immunological reagents and chemicals

A recombinant full-length HSV-2 186 GST-UL25 fusion protein was produced in *E. coli* strain BL21. Bacteria were lysed, and inclusion bodies were purified using a B-Per protein purification kit (Thermo Fisher Scientific, Ottawa, ON) according to the manufacturer’s instructions. Proteins in inclusion bodies were separated on preparative SDS-PAGE gels, the band corresponding to the glutathione *S*-transferase fusion excised and sent to Virusys (Taneytown, MD) to immunize a goat for pUL25 antiserum production. Goat polyclonal antiserum against HSV-2 pUL25 was used for western blotting at a dilution of 1:15000. Purified mouse anti-human IRF-3 (BD PharMingen, San Diego, CA) was used at a dilution of 1:100, and Alexa Fluor 568-conjugated donkey anti-mouse (Thermo Fisher Scientific, Ottawa, ON) was used at a dilution of 1:1000 for indirect immunofluorescence microscopy. Mouse monoclonal antibody against HSV ICP5 (Virusys, Taneytown, MD) was used for western blotting at a dilution of 1:500. Mouse monoclonal antibody against β-actin (Sigma, St. Louis, MO) was used for western blotting at dilution of 1:2000. Rabbit polyclonal antisera against HSV pUL16 (31), a gift from Dr. J. W. Wills (Pennsylvania State University), was used for western blotting at a dilution of 1:3000. Mouse monoclonal antibody against HSV-1 pUL25 (13), a gift from Dr. F. L. Homa (University of Pittsburgh), was used for western blotting at a dilution of 1:2000. Chicken polyclonal antisera against HSV pUL17 (62), a gift from Dr. J. D. Baines (Cornell University), was used for western blotting at a dilution of 1:2500. Rat polyclonal antisera against HSV pUL21 (28) was used for western blotting at a dilution of 1:600. Syto13 (Invitrogen, 1987320) was used for confocal microscopy at a final concentration of 37nM. AF 488 picolyl azide and 5-ethynyl-2’-deoxycytidine (EdC) (Click Chemistry Tools, Scottsdale, AZ) were used for click chemistry according to the manufacturer’s instructions.

### Transmission electron microscopy

Vero cells were plated on 100mm dishes one day prior to infection. Vero cells were infected at a multiplicity of infection (MOI) of 3 for 18 hours with WT strains, Δ21 mutants, or Δ21/Δ16 mutants. Infected cells were washed with PBS three times before fixing in 1.5 ml of 2.5% EM grade glutaraldehyde (Ted Pella, Redding, CA) in 0.1 M sodium cacodylate buffer (pH 7.4) for 60 minutes. Cells were collected by scraping into fixative and centrifugation at 300 x g for five minutes. Cell pellets were carefully enrobed in an equal volume of molten 5% low-melting temperature agarose (Lonza, Rockland, ME) and allowed to cool. Specimens in agarose were incubated in 2.5% glutaraldehyde in 0.1M sodium cacodylate buffer (pH 7.4) for 1.5 hours and post-fixed in 1% osmium tetroxide for one hour. The fixed cells in agarose were washed with distilled water three times and stained in 0.5% uranyl acetate overnight before dehydration in ascending grades of ethanol (30%-100%). Samples were transitioned from ethanol to infiltration with propylene oxide and embedded in Embed-812 hard resin (Electron Microscopy Sciences, Hatfield, PA). Blocks were sectioned at 50-60ριm and stained with uranyl acetate and Reynolds’ lead citrate. Images were collected using a Hitachi H-7000 transmission electron microscope. A-, B- and C-capsids were quantified in the nucleus (n = 9-12 micrographs per condition) and cytoplasm (n = 11-14 micrographs per condition), as well as in PEVs (n = 11-27 micrographs per condition).

### Evaluation of nuclear envelope integrity

To examine the localization of EGFP-JUMBO by microscopy, Vero cells growing on 35mm glass-bottom dishes (MatTek, Ashland, MA) were transfected using X-treme GENE HP DNA transfection reagent (Roche, Laval, QC) according to manufacturer’s instructions. At 24 hours post-transfection, cells were mock-infected or infected with HSV-1 F WT, or HSV-1 F Δ21. Cells were fixed with 4% paraformaldehyde in PBS and stained for pUL31 and DNA (Hoechst 33342) as described previously (27). Images were captured with an Olympus FV1000 laser scanning confocal microscope using a 60X (1.42 NA) oil immersion objective lens and FV10 ASW 4.01 software. Pixel intensities were measured using FluoView software version 4.01 and nuclear:cytoplasmic ratios of EGFP-JUMBO fluorescence intensity were quantified for n = 7 cells per condition. Composites of representative images were prepared using Adobe Photoshop software.

### Preparation of EdC labelled viruses

To incorporate EdC into viral genomes, T12 cells were seeded onto 150mm dishes and grown to confluence. Three days after confluency was reached cells were infected at an MOI of 0.01. At 4 hpi, 5-ethynyl-2’-deoxycytidine (EdC) was added directly to the medium at a final concentration of 1µM and fresh EdC was added every 24 hours until complete cytopathic effect was observed. Cells were scraped into the medium and virus stocks were prepared as described previously (63). Virus stocks were cleared of residual EdC using a PD-10 desalting column (GE Healthcare, Mississauga, ON) utilizing the manufacturer’s instructions. Desalted virus preparations were aliquoted and stored at -80°C.

### Analysis of extracellular virions

Infected cell culture supernatants, containing extracellular virions, were cleared of cellular debris by three consecutive centrifugations at 1,000 x g for 10 minutes at 4°C. After the final centrifugation, supernatants were filtered using a 0.45µm filter and 1.2mL of the supernatants were transferred into polycarbonate TLA 100.3 centrifuge tubes. Extracellular virions were then treated with 2µL of benzonase (250U/µL) (Santa Cruz Biotechnology, Dallas, TX) and incubated at RT for 10 minutes. Extracellular virions were pelleted in a TLA 100.3 rotor at 25,000 rpm for 36 minutes at 4°C using a Beckman Optima Max-XP ultracentrifuge. Supernatants were removed and virus pellets resuspended in 600µL of cold PBS. The extracellular virions were then transferred to 1.5mL centrifuge tubes and virions dispersed by gentle sonication in a chilled cup-horn sonicator. Syto13 (Thermo Fisher Scientific, Ottawa, ON) (8mM in DMSO) was added to samples at a final dilution of 1:135,000 that were then incubated at 4°C overnight. Syto13 labelled virions were then plated on poly-L-lysine coated 35mm glass-bottom dishes and incubated on ice for 30 minutes at 4°C. Extracellular virions were imaged through a 60X (1.42 NA) oil immersion objective using an Olympus FV1000 confocal laser scanning microscope and FV10 ASW 4.01 software. Capsids in these images were then counted utilizing Image Pro Plus software, version 6.3.0.512. Non-genome containing capsids (A- or B-capsids) were identified by a red signal, while genome-containing capsids (C-capsids) were identified by a yellow signal indicating the colocalization between genomic DNA (green) and capsid (red) signals. Aggregates of yellow and red signals as well as lone green signals were not counted. A total of n = 30 images per condition over three biological replicates were analyzed.

### Nuclear translocation of IRF-3

WT viruses and their corresponding Δ21 mutants were used to infect T12 cells on ice at an MOI of 3. After one hour, the inoculum was replaced with complete medium containing 50µg/mL of cycloheximide. At 0, 2, 4 and 6 hpi, cells were washed three times with PBS and fixed by adding 4% formaldehyde in PBS for 15 minutes at room temperature. Cells were permeabilized and stained using a mouse monoclonal IRF-3 antibody as described previously (27). As a positive control for IRF-3 nuclear translocation, T12 cells were transfected with 2µg poly I:C (InvivoGen, San Diego, CA) using X-treme GENE HP DNA transfection reagent (Roche, Laval, QC) following the manufacturer’s instructions. At 6 hours post-transfection, cells were fixed and stained for IRF-3.

### Capsid isolation

Capsid preparation was performed similarly to previously published methods (57). Five 150mm dishes of confluent Vero cells were infected with each HSV WT, or their corresponding Δ21, strains. When complete CPE was reached, infected cells were harvested and centrifuged at 1,000 x g for 10 minutes at 4°C. Supernatants were discarded and cell pellets were resuspended in 50mL of cold PBS and pelleted at 1,000 x g for 10 minutes at 4°C. The supernatants were discarded, and cells were lysed by resuspension in 10mL cold NP-40 lysis buffer (150mM NaCl, 10mM Tris pH 7.2, 1% NP-40, 5mM DTT, 2mM MgCl_2_) containing protease inhibitors (Roche, Laval, QC) on ice for 30 minutes. The nuclei were then pelleted at 1,000 x g for 10 minutes at 4°C, supernatants were discarded, and nuclei were resuspended in 3mL of TNE buffer (150mM NaCl, 20mM Tris pH 7.5, 1mM EDTA). Nuclei were then broken by successive passages through 18-, 22- and 25-gauge syringe needles. Nuclear lysates were treated with 1µL of benzonase (250U/µL), incubated for 15 minutes at RT and then clarified by centrifugation at 3,000 x g for 10 minutes at 4°C. Clarified nuclear lysates were layered onto a 2mL 35% (w/v) sucrose cushion prepared in TNE and centrifuged at 35,000 rpm for 32 minutes at 4°C in a Beckman SW55 rotor. Pellets containing nucleocapsids were resuspended in 250µL of TNE buffer and sonicated briefly using a chilled cup-horn sonicator. Capsids were then layered onto 10mL 20% to 50% linear sucrose gradients prepared using a Gradient Master (BioComp, Fredericton, NB) and centrifuged in a Beckman SW41 rotor at 25,000 rpm for 1 hour. After centrifugation, distinct light-scattering bands representing A-, B-, and C-capsids were evident. For capsids to be analyzed by western blotting, gradients were fractioned from the top of the tube into 1mL fractions. For capsids analyzed by mass spectrometry, capsids were isolated from the gradient by side wall puncture using a 16 ½-gauge needle. Samples collected from the gradient were diluted with 0.5 volumes of TNE buffer and capsids pelleted at 26,000 rpm for 30 minutes at 4°C in a Beckman MLA130 rotor. Supernatants were discarded and capsid pellets were resuspended in 35µL of 1X SDS-PAGE loading buffer and stored at -20°C until analyzed by western blotting or mass spectrometry.

### Mass spectrometry of nuclear capsids

HSV-1 KOS WT and Δ21 nuclear A-, B-, and C-capsids were electrophoresed 0.5 cm into a 9% SDS-PAGE gel. Protein bands from each sample were visualized by staining with SimplyBlue (Thermo Fisher Scientific, Ottawa, ON). Proteins were excised in a 0.5cm by 0.5cm slab of acrylamide. Slabs were placed in 2mL microcentrifuge tubes containing 500µL of 1% acetic acid and sent on wet ice to the Southern Alberta Mass Spectrometry Facility for liquid chromatography tandem mass spectrometry.Proteins identified by mass spectrometry that met the following criteria were included in the analysis: a >95% protein and peptide threshold, a minimum number of ≥3 unique peptides identified, and by removing common contaminants (e.g., keratins). To ensure direct comparisons could be made between WT and Δ21 samples, Δ21 A-, B-, and C-capsids were normalized to respective WT capsids using total spectrum counts for VP5. Three biological replicates were analyzed for each strain.

### Western blot analysis

Proteins were electrophoresed through 8% or 10% SDS-PAGE gels. Proteins were transferred onto polyvinylidene fluoride (PVDF) membranes and blocked for 1 hour in Intercept tris-buffer saline (TBS) blocking buffer (LI-COR, Lincoln, NE). Membranes were probed with appropriate dilutions of antibodies (VP5, pUL25, pUL17, pUL21, or pUL16) overnight at 4°C. Membranes were washed three times with Tris-buffered saline (10mM Tris pH 8.0, 150mM NaCl) containing 0.1% Tween20 (TBST) for seven minutes and then appropriate dilutions of secondary antibodies were added to the membranes and incubated at RT for one hour. Membranes were then washed three times with TBST for seven minutes and finally with TBS before imaging on an Odyssey CLx imaging system (LI-COR, Lincoln, NE). Images and signal intensities of viral proteins were obtained using Image Studio Lite software version 5.2.5. VP5 signals were used to normalize WT and Δ21 capsid numbers. Normalized signal intensity for pUL21, pUL16, pUL25 and pUL17 in Δ21 A-, B-, and C-capsid samples were directly compared to corresponding WT capsids samples.

## Acknowledgements

This work was supported by the Canadian Institutes of Health Research operating grant 93804, Natural Sciences and Engineering Research Council of Canada Discovery Grant 418719 and Canada Foundation for Innovation award 16389 to BWB. The funders had no role in study design, data collection and interpretation, or the decision to submit the work for publication. We thank Drs. Fred Homa (University of Pittsburgh), Joel Baines (Cornell University), John Wills (Pennsylvania State University) for kindly providing antisera, Dr. Wade Bresnahan (University of Minnesota) for providing T12 cells, Dr. Renée Finnen for providing pRF52, and Dr. Greg Smith (Northwestern University) for providing the VP26-mRFP1 HSV-1 strain. We are grateful to Drs. Renée Finnen, Jie Gao and Xiahou Yan (Queen’s University) for assistance with electron microscopy, and Dr. Laurent Brechenmacher (Southern Alberta Mass Spectrometry Centre) for assistance with mass spectrometry. We are indebted to our colleagues Renée Finnen and Safara Holder for helpful discussions and comments on the manuscript.

